# Early urine proteome changes in an implanted bone cancer rat model

**DOI:** 10.1101/613125

**Authors:** Ting Wang, Lujun Li, Weiwei Qin, Yuhang Huan, Youhe Gao

## Abstract

In this study, Walker 256 cells were implanted into rat tibiae. Urine samples were then collected on days 3, 5, 7, and 13 and were analyzed by liquid chromatography-tandem mass spectrometry (LC-MS/MS). With label-free quantification, 25 proteins were found to change significantly in the urine of the tumor group mice compared with the proteins in the urine of the control group mice; this was even the case when there were no significant lesions identified in the Computed Tomography(CT) examination. Among these differentially proteins, 7 were reported to be associated with tumor bone metastasis. GO analysis shows that the differential proteins on day 3 were involved in several responses, including the acute phase response, the adaptive immune response and the innate immune response. The differentially proteins on day 7 were involved in the mineral absorption pathway. The differentially proteins on day 13 were involved in vitamin D binding and calcium ion binding. These processes may be associated with bone metastasis. Our results demonstrate that urine could sensitively reflect the changes in the early stage of implanted bone cancer; this provides valuable clues for future studies of urine biomarkers for tumor bone metastasis.

## Introduction

Biomarkers are the measurable changes associated with a physiological or pathophysiological process. When the body is stimulated by physiological or pathological factors, most of the changes are excreted in the blood for a short period of time because the blood has a strict steady-state regulation. As an excrement, urine can accumulate more changes. Additionally, change is the essence of disease markers. Therefore, many changes in the body can be reflected in the urine. The urine can reflect the changes in multiple organs of the body well, and it is more sensitive to diseases or to physiological changes. Therefore, it is easier to find biomarkers in the urine [1-3]. Although urine is an ideal source of biomarkers, the composition of urine is very complex. This is especially true in human urine samples; in addition to the markers produced by the disease, there are many other interfering factors. This makes it difficult to directly use human clinical urine samples to find disease markers. It is not easy to detect markers in the early stages of the disease. Therefore, there is an urgent need to find an easier way to find these markers. We believe that by establishing animal models of disease, biomarkers can be screened; this can remove the influence of other complex factors on the markers and can screen out potential biomarkers at an earlier stage and more easily [4].

Our previous laboratory studies have also found that urine can respond to changes in the body due to disease in an early and sensitive manner, and we have found that it can reflect a number of systemic diseases. In terms of kidney disease, the urinary proteins of two different glomerular injuries changed significantly before the histopathological changes in the kidney were observed [5]. The urinary proteins changed significantly in the renal interstitial fibrosis model constructed by unilateral ureteral ligation [6]. In the model of focal segmental glomerulosclerosis induced by doxorubicin, the urinary proteins were continuously changing with disease progression [7]. There are other models of organ diseases as follows: bleomycin-induced pulmonary fibrosis model [8], experimental autoimmune myocarditis model [9], rat model of bacterial meningitis [10], glioblastoma animal model [11], tumor model of the subcutaneous implantation of Walker 256 cells [12], rat model of chronic pancreatitis induced by diethyldithiocarbamate [13], rat model of NuTu-19 lung metastasis [14], Walker 256 intracerebral tumor model [15], and Walker 256 lung metastasis rat model [16]. The urinary proteome changes significantly before the clinical signs and histopathological changes have occurred. In summary, early laboratory results can show that the urine can reflect the changes caused by the disease in an early and sensitive way.

Bone is a common metastatic site for malignant tumors and is second only to the lungs and liver. The data indicate that 75%-90% of patients with advanced tumors develop bone metastases and experience significant cancer pain [17]. Tumor bone metastasis affects the quality of life of patients and causes patients to suffer from anemia, fractures, paraplegia, hypercalcemia, pain and cachexia; in addition, it increases patient mortality [18]. Therefore, the early diagnosis and treatment of tumor bone metastasis is of great significance to improve the patient’s quality of life and to prolong the survival rate. A needle biopsy is the “gold standard” for diagnosing bone metastases. However, due to its invasive examination, clinical compliance is low. Imaging examination plays an irreplaceable role in the screening and early diagnosis of bone metastases. The imaging methods for diagnosing bone metastases mainly include X-ray, CT, MRI, whole body bone imaging (WBS), SPECT, and PET/CT. However, all of the above methods have defects, and their ability to achieve an early diagnosis is not ideal. Bone metastasis is usually diagnosed when it is discovered [19-20]. At the time of bone metastasis, the changes in the biochemical markers of bone metabolism were significantly earlier than the morphological changes found in the imaging studies. Researchers began to pay attention to bone metabolism indicators and search for disease markers [21]. We focus on urine, which is more ideal to screen tumor bone metastasis markers.

The Walker 256 cells used in this experiment were found in the mammary glands of pregnant rats (*Rattus norvegicus*) and are considered a type of carcinosarcoma [22-23]. It is the most widely used transplantable tumor in experimental research. The cells can cause significant bone resorption and bone destruction at the site of rat bone implantation this is consistent with the phenotype of bone metastasis in breast cancer patients [24]. In this experiment, we implanted Walker 256 cells into the left tibia of the rat. Urine was collected for analysis in the early, middle, and late stages of tumor growth. We also used this cell type because our laboratory has used this cell for other tumor studies. The purpose of the experiment was also to explore the changes caused by the same tumor cells in different parts of the body. We then explored whether these changes can be reflected in the urine.

This experiment established an implanted bone cancer rat model. The urine samples of the model rats were analyzed by proteomics analysis, which identified differential proteins that can reflect the state of the model early.

## 1. Materials and methods

### 1.1 Establishment of the bone metastasis model

Adult SPF male Sprague-Dawley rats (180-200 g) were provided by Beijing Vital River Laboratory Animal Technology Co., Ltd. The animal license is SCXK (Beijing) 2016-0006. Animal experiments were reviewed and approved. Rats are housed in a constant temperature, well ventilated environment with adequate food and water. Walker 256 cancer cells were passaged in ascites using SD pups. The young rats weighed 60-80 g and were male rats.

Walker 256 cancer cells were passaged in ascites, and the method was based on previous studies [25].1.1.2 Preparation of the rat model using the Walker 256 cell line. The experimental method was improved based on the preceding modeling method [26]. Briefly, the rats were anesthetized by intraperitoneal injection of 2% sodium pentobarbital (30 mg/kg). A 10 ml syringe needle was slowly rotated on the tibia. A cell suspension of 100 cells was injected along the pinhole with a 10 μL microinjector. The rats in the control group were injected only with saline.

### 1.2 CT examination of the tibia of the rat model

We used SIEMENS Inveon preclinical (small animal) PET-CT to perform CT imaging on the left leg of the rat and to observe the progress of bone metastasis in the left leg of the rat.

### 1.3 Urine sample preparation

After modeling, urine samples from the rats of the control group and the tumor group were collected using a rat metabolic cage, and the urine samples were stored in a freezer at −80 °C. The process of extracting the urinary proteins was as follows: The nontreated rat urine was frozen in a tube, thawed in a 37 °C water bath, and centrifuged at 4 °C at 3000 × g for 30 min. The supernatant was centrifuged at 4 °C at 12 000 × g for 30 min to remove large cell debris. The protein was precipitated using 3 times the volume (or more) of prechilled ethanol at –4 °C overnight. The precipitation process was described previously [27]; after the urine was precipitated overnight, it was centrifuged to retain the protein precipitation, and the urinary protein was dissolved by the lysate (8 mol/L urea, 2 mol/L thiourea, 25 mmol/L dithiothreitol and 50 mmol/L Tris). The protein solution was centrifuged to obtain a supernatant, and the protein concentration was determined by the Bradford method; the supernatant was used to run an SDS-PAGE gel and proteolysis.

The six rats in the control group and the tumor group were subjected to mass spectrometry by membrane digestion. The process of enzymatic cleavage was performed as described previously [28]: the protein was loaded onto a 10 kDa filter (Pall, Port Washington, NY, USA); the urinary proteins were washed sequentially with UA (8 mol/L urea plus 0.1 mol/L Tris-HCl, pH 8.5) and 50 mmol/L NH4HCO3. Then, 5 mmol/L DTT and 50 mmol/L IAA were added to the membrane system. After adding trypsin (1:50) for an overnight digestion, the peptides were collected. The peptides were desalted by an HLB column (Waters, Milford, MA) and were then drained by a vacuum pump (Thermo Fisher Scientific, Bremen, Germany). The peptides were acidified with 0.1% formic acid and were loaded onto a reversed-phase microcapillary column using a Thermal EASY-n LC1200 chromatography system. Data were acquired by mass spectrometry using a Thermo Orbitrap Fusion Lumos mass spectrometer (Thermo Fisher Scientific, Bremen, Germany).

### 1.4 Liquid chromatography-mass spectrometry/mass spectrometry analysis

The digested peptides were dissolved in 0.1% formic acid and were loaded onto a trap column (75 μm × 2 cm, 3 μm, C18, 100 Å). The eluent was transferred to a reverse-phase analytical column (50 μm × 150 mm, 2 μm, C18, 100 Å) by a Thermo EASY-nLC 1200 high-performance liquid chromatography system. The peptides were analyzed using a Fusion Lumos mass spectrometer (Thermo Fisher Scientific). The Fusion Lumos was operated on data-dependent acquisition mode. The survey mass spectrometry (MS) scans were acquired in the Orbitrap using a 350–1550 m/z range with a resolution set to 120,000. The most intense ions per survey scan (top speed mode) were selected for collision-induced dissociation fragmentation, and the resulting fragments were analyzed in Orbitrap. Dynamic exclusion was employed with a 30 s window to prevent the repetitive selection of the same peptide. The normalized collision energy for HCD-MS2 experiments was set to 30%. Two technical replicate analyses were performed for each sample.

### 1.5 Database searching

Mascot software (2.4.0) was used for database retrieval. The details are as follows: 1) Raw files collected by the mass spectrometer were extracted as secondary spectral peaks and were converted into MGF format files. 2) All MGF format files were imported into Mascot software. 3) The parameters of the database search were set, and the Swiss-Prot database was used (data up to June 3, 2017). Other retrieval conditions were as follows: rat species; tryptase digestion, which allowed two tryptase leak sites at most; urea methylation fixed with cysteine, which could be modified with N-terminal acetylation and oxidation modification; mother ion mass accuracy was 0.02 Da, and the daughter ion mass accuracy was 10 ppm. 4) Screening conditions after searching: peptide level FDR < 1%, protein level FDR < 1%, ion score > 30. Finally, the search results were exported in Xml format for the subsequent quantitative analysis.

### 1.6 Label-free quantification

Quantitative analysis is based on MS1. The acquired spectra were loaded into Progenesis software (version 4.1, Nonlinear, Newcastle upon Tyne, UK) for label-free quantification, as described previously (Hauck et al., 2010). Briefly, features with only one charge or with more than five charges were excluded from the analyses. For further quantitation, all peptides (with Mascot score >30 and P<0.01) of an identified protein were included. Proteins identified by at least one peptide were retained. The MS/MS spectra exported from Progenesis software were processed with Mascot software (version 2.4.1, Matrix Science, UK) in the Swiss-Prot database (data 06/03/2017). The search parameters were set as described before.

Quantitative analysis is based on the number of MS2. Scaffold software and MASCOT 2.4.0 software were used to screen and integrate the search results and to quantify the number of spectra. 1) The DAT files exported from Mascot software after searching the database are imported into Scaffold software one-by-one. 2) The search parameters were set as described in the database searching methods section 3) The retrieval conditions were as follows: the identification conditions of the peptide segments were reliability (> 90.0%) and FDR (< 0.1%); the protein identification conditions were reliability (> 95.0%) and that each protein contained at least two identified peptide segments.

### 1.7 Statistical analysis

The identified proteins were screened. The screening conditions were as follows: fold change of protein abundance > 2 times; p-value < 0.05. Each normalized abundance in the experimental group was greater or less than each normalized abundance in the control group. Then, the UniProt database was used to identify the corresponding human homologous proteins to the rat proteins.

### 1.8 Gene ontology analysis

The differential proteins were analyzed using the online database DAVID (https://david.ncifcrf.gov/) for GO analysis. The results of the analysis include the four following aspects: molecular function, biological processes, cellular components, and pathways. The above results are shown using graphs and tables.

## 2. Results and analysis

### 2.1 Weight changes with disease

As the disease progressed, the body weights of the experimental group mice increased more slowly than those of the control group mice on the seventh day after tumor cell implantation (Fig. 1). On the tenth day, the weight dropped. Both the amount of ingested food and water were significantly reduced. The activity of rats in the experimental group decreased, which may be caused by bone pain.

**Figure 1:**
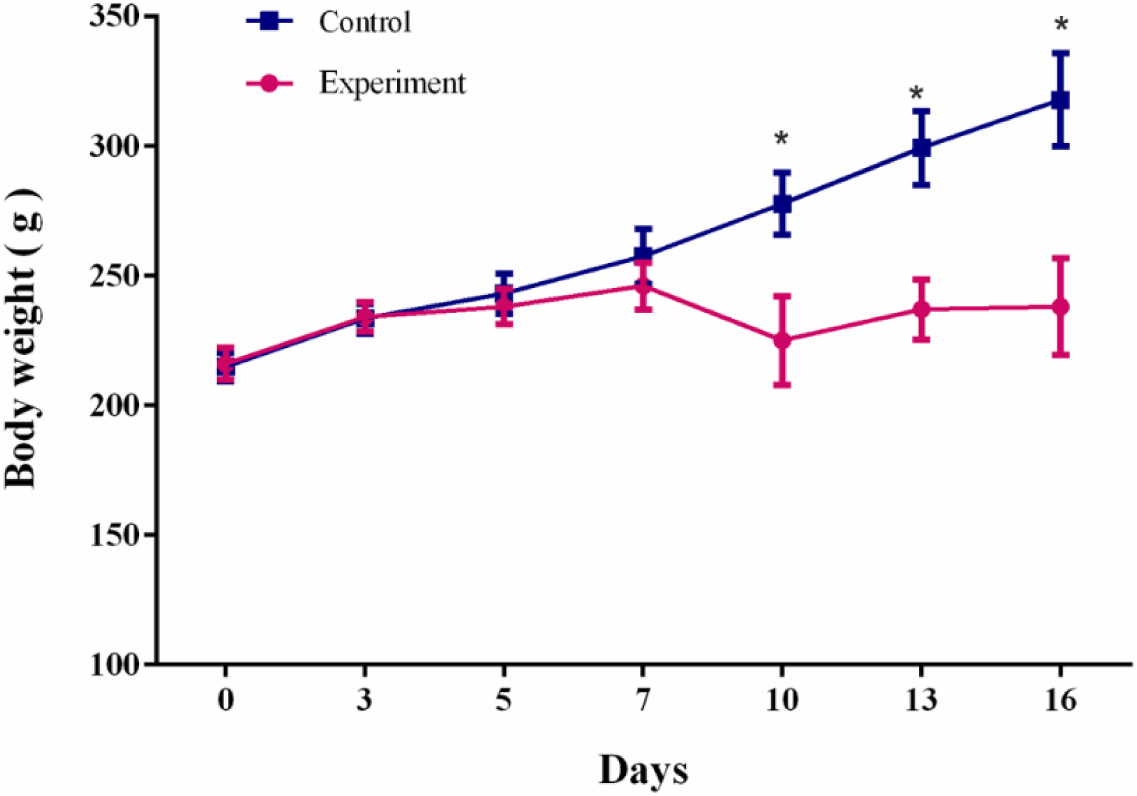
Comparison of body weight between Walker 256 model rats and control rats (* p<0.01).

### 2.2 CT results of rat tibia

It can be seen from the CT results (Fig. 2) that as the disease progresses, the tumor group rats have tumor cells that aggregate and grow near the pinhole on the fifth day compared with those of the control rats. On the tenth day, the tumor cells invaded the bone more deeply, and the bone density decreased significantly. The CT results are basically consistent with the changes in body weight, indicating that this is a successful model.

**Figure 2:**
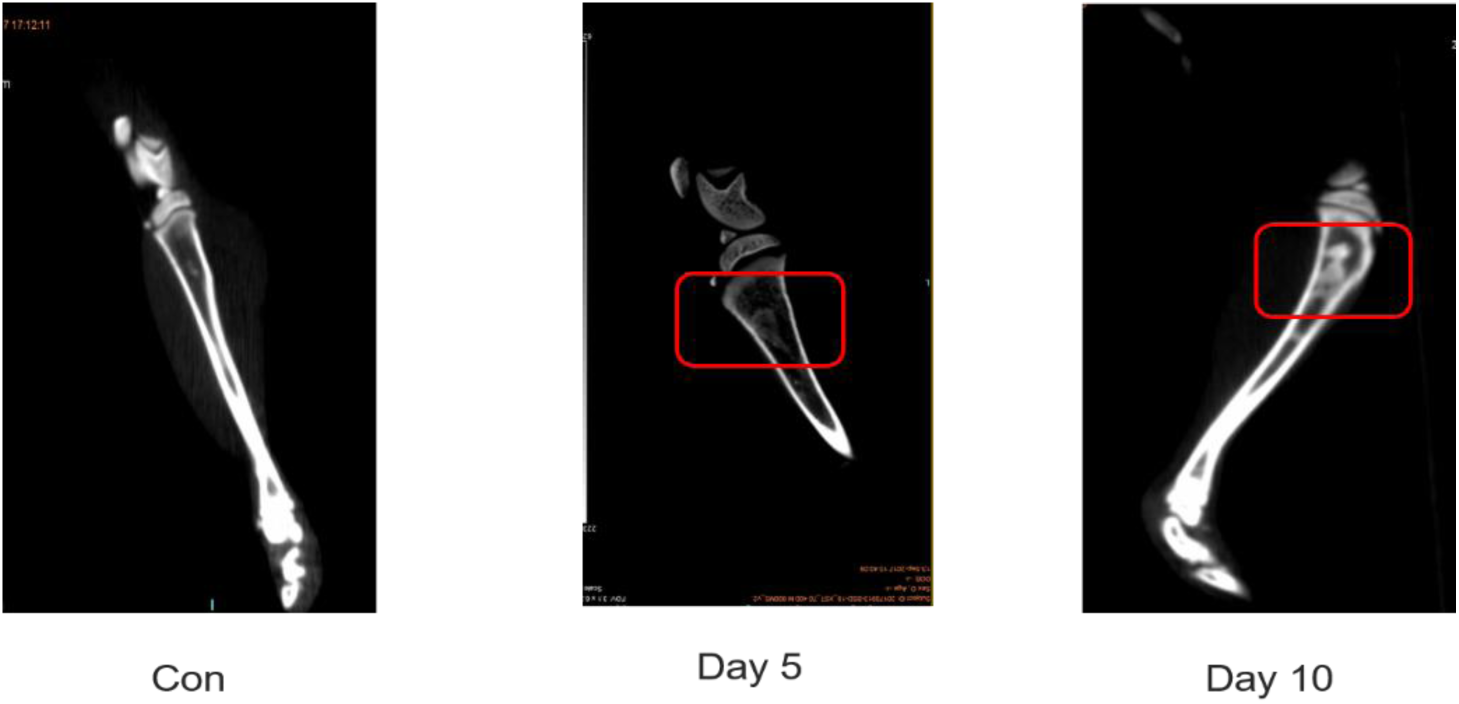
CT results of rat tibiae. (Con: Normal control rat; Day 5: Walker 256 cells implanted on day 5; Day 10: Walker 256 cells implanted on day 10; The part marked in red: Tumor cell aggregation and growth near the pinhole).

### 2.3 Changes in the urinary proteins with disease progression (SDS-PAGE)

The urinary proteins on Day 3, Day 5, Day 7 and Day 13 of the control group rats and the tumor group rats were analyzed by SDS-PAGE gel electrophoresis. It was found that the expression of the urinary protein at 25 kDa increased as the degree of tumor invasion increased. The results of the changes in several rats were consistent (Fig. 3). According to the experimental results, the protein at 25 kDa may be NGAL (Neutrophil gelatinase-associated lipocalin), NKG2D (NKG2-D type II integral membrane protein). These proteins were highly abundant, the fold change increased with time, and the molecular weight was 25 kDa.

**Figure 3:**
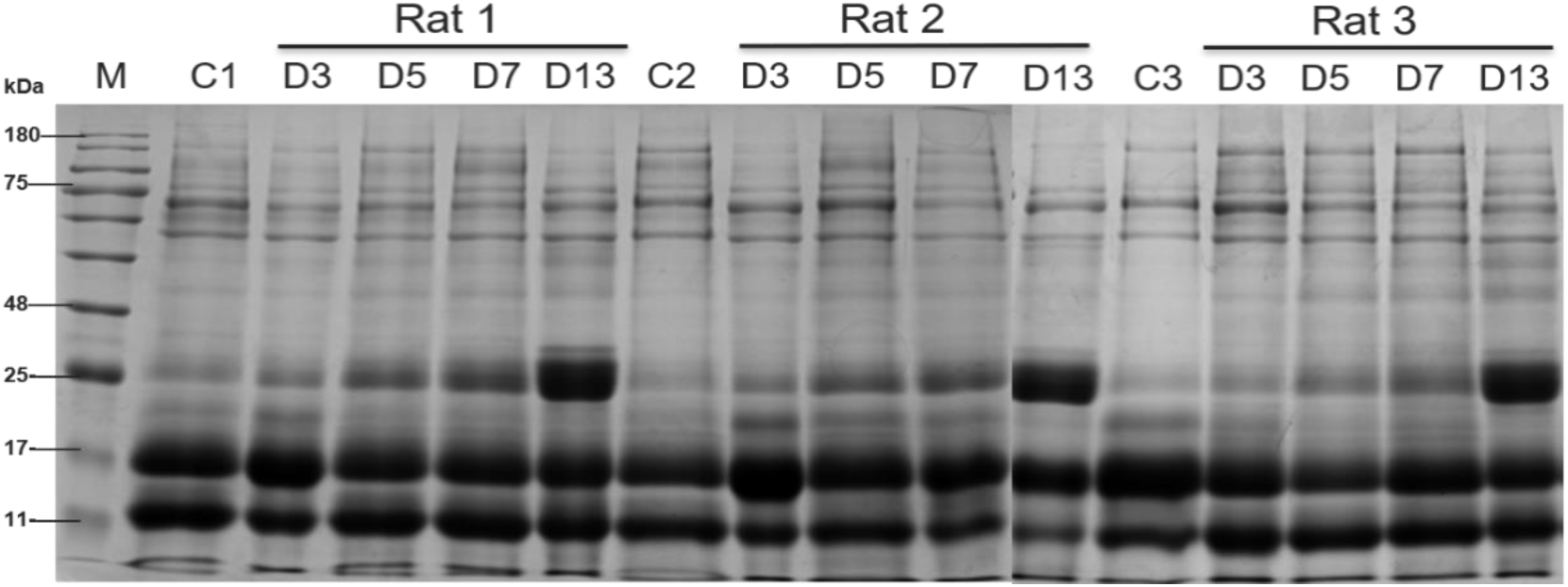
Analysis of urinary protein SDS-PAGE results in model rats. (M: marker; C1, C2, C3: control group 1, 2, 3; D3, D5, D7, D13: Walker 256 cells implanted on day 3, 5, 7, 13; Rat 1, 2, 3: Model group rat 1, 2, 3).

### 2.4 Identification of urinary proteins

After LC-MS/MS identification, a total of 900 urinary proteins were identified. Urinary proteins that meet the following criteria can be screened for differential proteins: 1) P < 0.05; 2) fold change ≥ 2 or ≤ 0.5; 3) Normalized abundance for each sample in the elevated group is greater than that of each sample of the control group with normalized abundance. The descending group is the opposite.

Tumor group rats were screened for 27 differential proteins on Day 3, of which, 25 proteins had human homologous proteins (Table 1). Of the 25 proteins, 23 were upregulated and 2 were downregulated. Seventeen different proteins were screened on Day 5, of which, 13 proteins had human homologous proteins (Table 1). Ten of the 13 proteins were upregulated, and 3 were downregulated. Twenty-four differentially proteins were screened on Day 7, of which, 20 differentially proteins had human homologous proteins (Table 1). Seventeen of the 20 proteins were upregulated, and 3 were downregulated. Thirty different proteins were screened on Day 13, of which, 27 differential proteins had human homologous proteins (Table 1). Seven of the 27 proteins were upregulated, and 20 were downregulated.

**Table 1:**
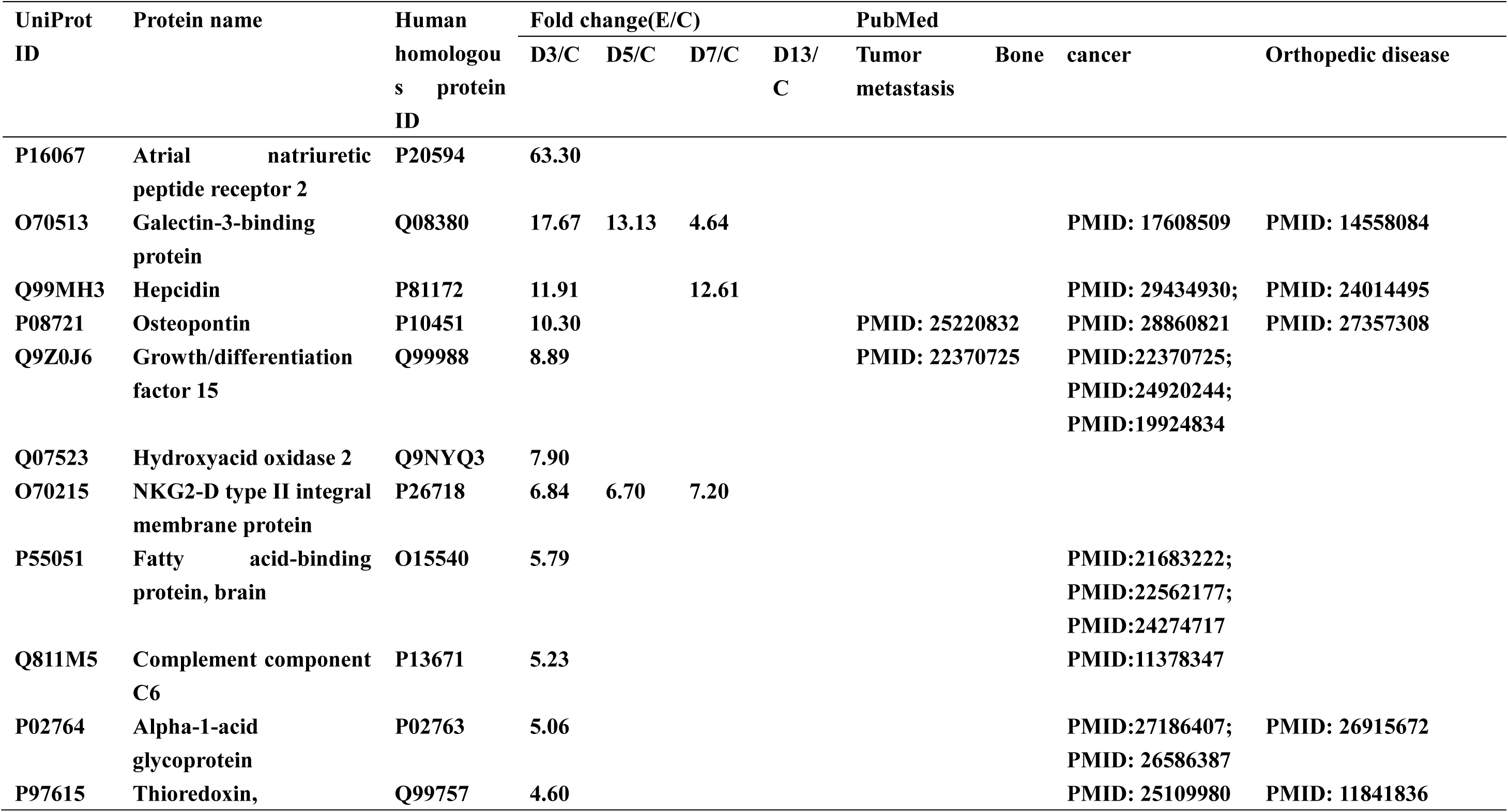
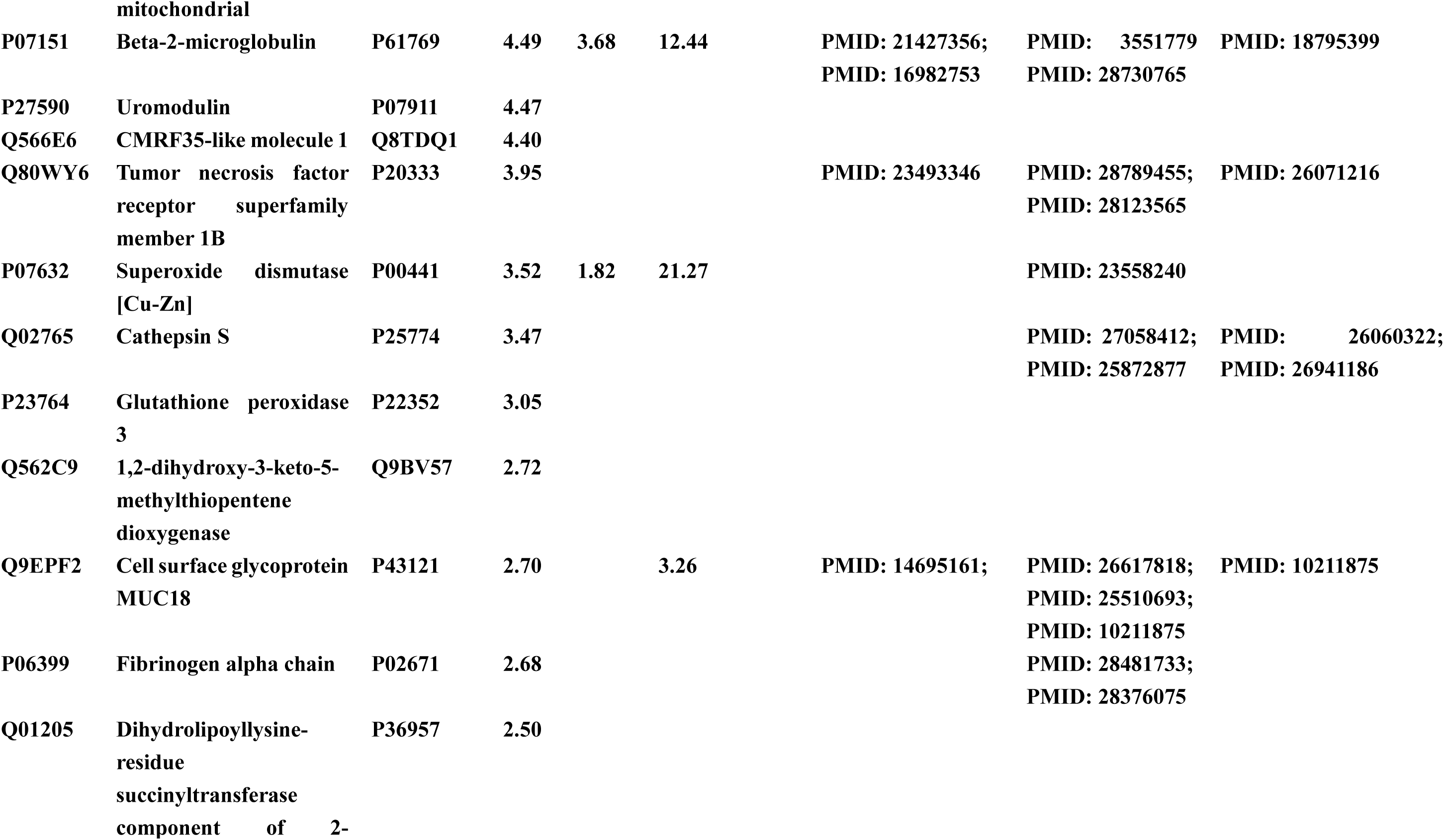
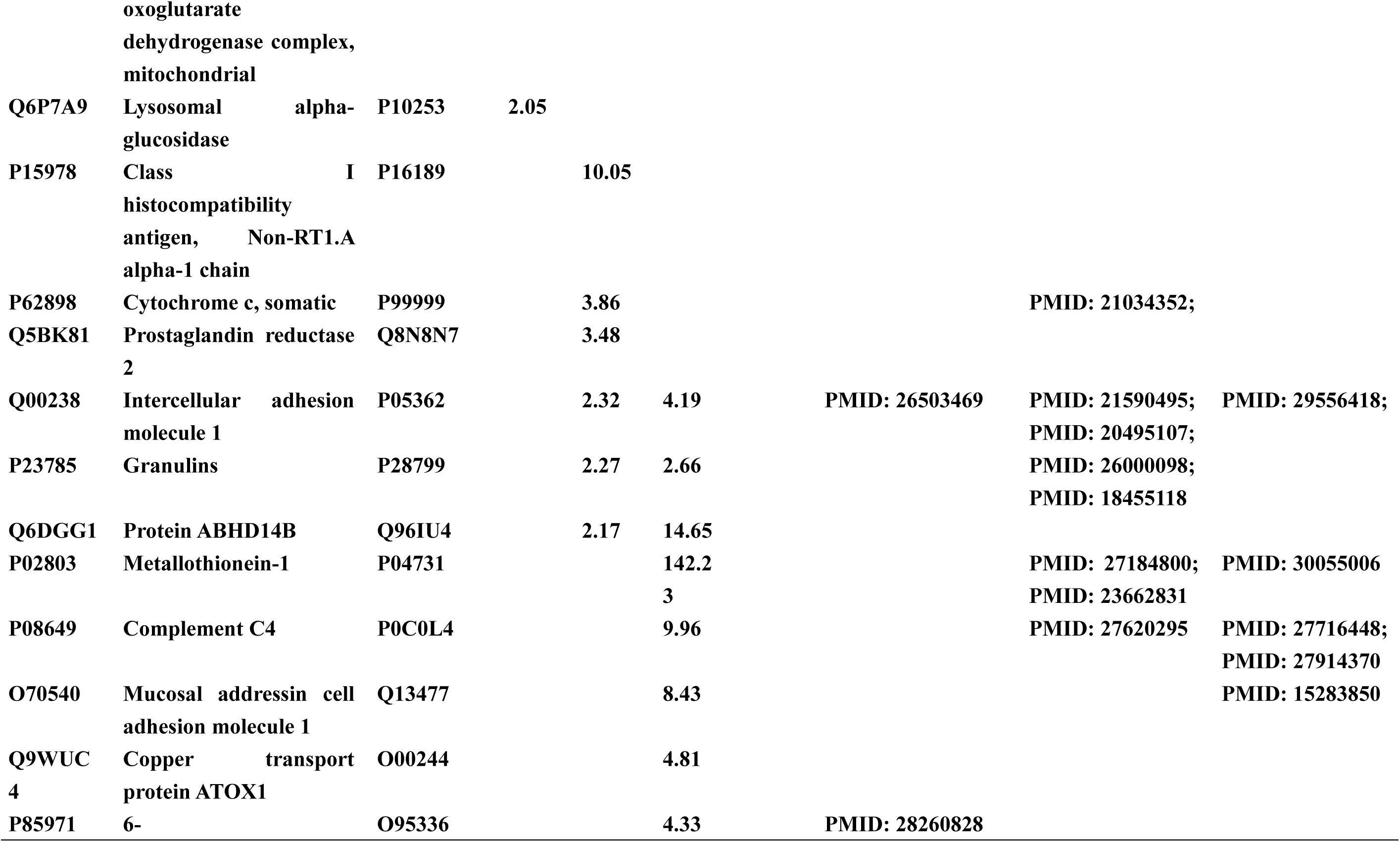
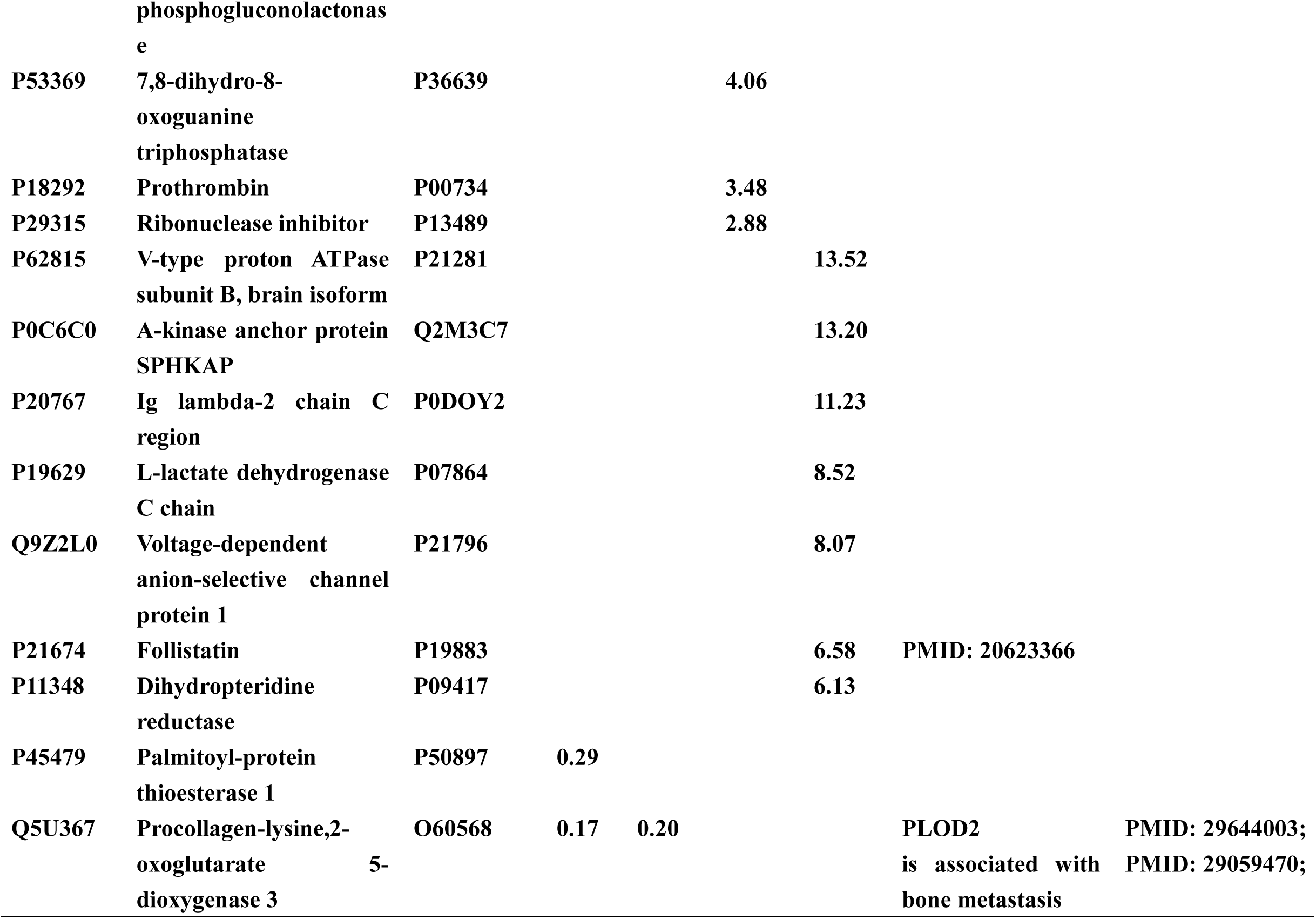
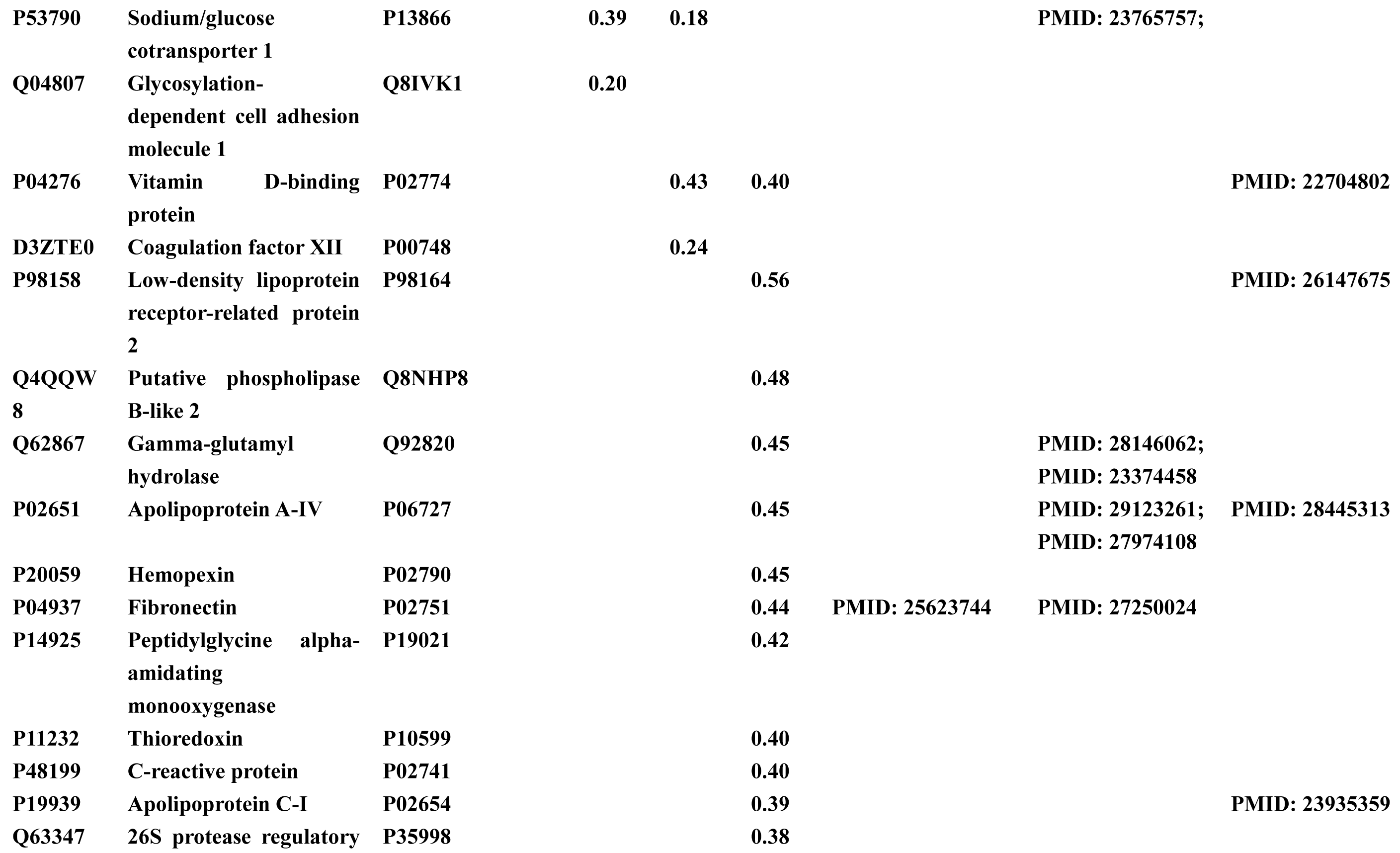
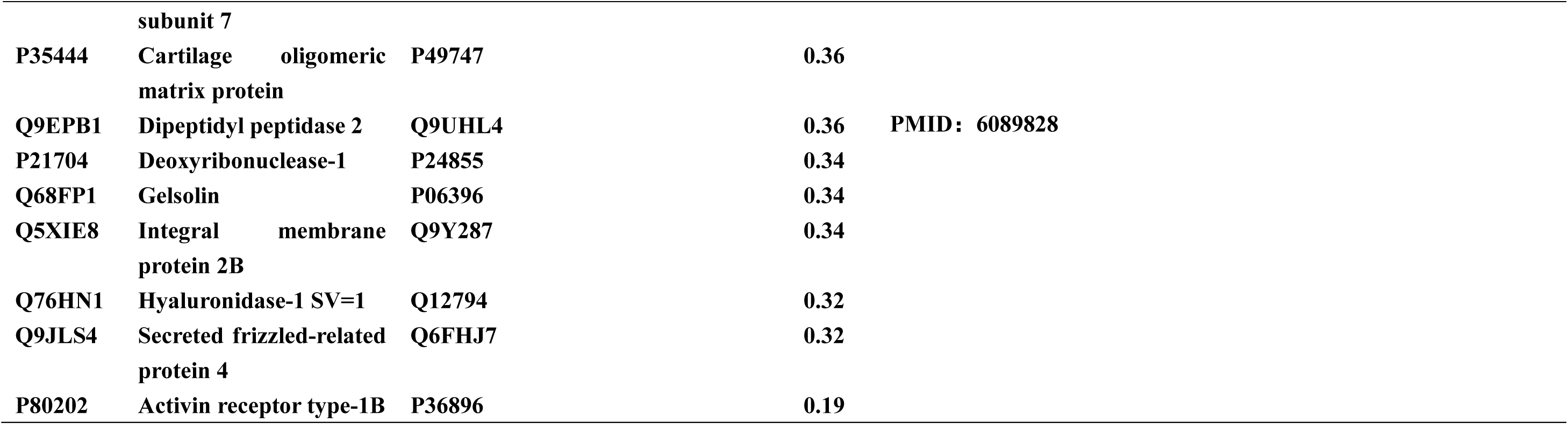
Differential protein statistics at various time points in the rat model of tumor bone metastasis and their association with related diseases.

The above differential proteins were subjected to Venn diagram statistics. The changes in the differential proteins with different time points gave the following results (Fig. 4). As seen from the results below, a total of 69 differential proteins were identified, and no protein changed with time. There are separately varying differential proteins at each time point. There are four differential proteins that have been changing on Days 3, 5, and 7.

**Figure 4:**
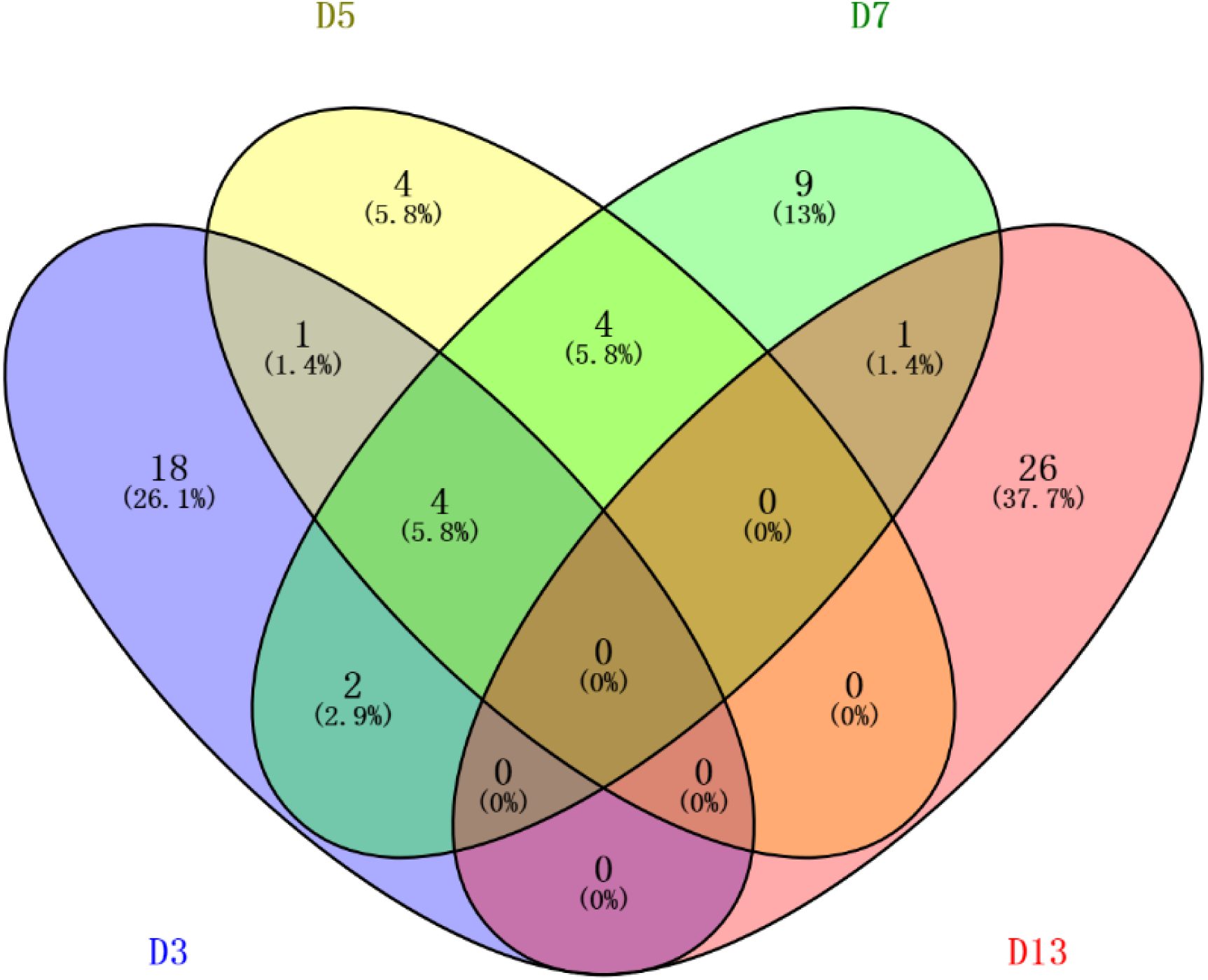
Differential protein changes at various time points compared to the control group. (D3, D5, D7, and D13: Walker 256 cells implanted on day 3, 5, 7, and 13).

In the screening and statistics of differential proteins, we focused on finding the relationship between the differential proteins and the tumor bone metastases, bone-related diseases, and tumors in previous studies. The statistics are shown in Table 1 below.

By searching many studies, we found that many of the different proteins obtained were related to the above three processes or were a direct marker (Table 1). For example, in the differential protein on the third day after tumor implantation, some proteins were associated with tumor bone metastasis as follows:

a. Osteopontin, an important factor in the bone metastasis of lung cancer, prostate cancer and breast cancer, is a marker of bone metastasis (PMID: 25220832), is associated with breast cancer (PMID: 28860821), is involved in bone remodeling and promotes disease progression in osteoarthritis (PMID: 27357308).
b. Growth/differentiation factor 15 regulates osteoclast differentiation and can be used to treat prostate cancer bone metastasis (22370725), which is associated with prostate cancer progression (22370725), and it is an effective biomarker for patients with metastatic colorectal cancer (29400662).
c. Tumor necrosis factor receptor superfamily member 1B plays an important role in bone metastasis (PMID: 23493346) and is associated with rheumatoid arthritis (PMID: 26071216), which plays an important role in the development and prognosis of breast cancer (PMID: 28789455). This protein is also a new target for colorectal cancer treatment (PMID: 28123565).
d. Cell surface glycoprotein MUC18 plays an important role in the metastasis of osteosarcoma (PMID: 14695161), which is an independent prognostic factor for renal cell carcinoma (PMID: 26617818) and a marker of the malignant potential of ovarian cancer. It is overexpressed in malignant melanoma cells, melanoma cells, and prostate cancer cells (PMID: 25510693), and it is a marker of the progression and metastasis of human melanoma (PMID: 10211875).
e. Procollagen-lysine, 2-oxoglutarate 5-dioxygenase 3 is a marker of precancerous lesions in early liver cancer (PMID: 29059470). PLOD2 in this family of proteins promotes bone metastasis and is associated with osteoarthritis.

The vast majority of differential proteins on the third day are associated with various tumors and are markers of numerous tumors. It can be seen from the above results that in the early stage of the tumor of the rat model (early stage of tumor cell implantation, and no significant change in CT and body weight), the changes in the body at the early stage of the disease can be seen from the urine.

On the fifth day, the continuously changing protein was removed, and the differential protein was associated with the disease. Intercellular adhesion molecule 1 is a factor in the bone metastasis of breast cancer and is associated with the aggressiveness of breast cancer (PMID: 21590495). It is associated with rheumatoid arthritis disease activity (PMID: 29556418) and can also aid in the diagnosis of lung cancer (PMID: 20495107). Other differential proteins at this time point are also mostly associated with tumors and even exist as markers.

The Galectin-3-binding protein, which is continuously changed in D3, D5, and D7, is a prominent marker of rheumatoid arthritis disease activity (PMID: 14558084) and is considered to be a tumor marker of breast cancer, prostate, colon and pancreatic cancer (PMID: 17608509). Beta-2-microglobulin is a growth-promoting factor for the bone metastasis of prostate cancer and is closely related to metastasis (PMID: 16982753). It is a tumor marker for nasopharyngeal carcinoma (PMID: 3551779) and is a predictor of breast cancer survival (PMID: 28730765). It was also found to be involved in the disease process of osteoarthritis (PMID: 18795399). On the seventh and thirteenth days (when CT found lesions), many differential proteins were also found to be associated with tumor bone metastasis.

As seen from the above analysis, among the differential proteins we screened, many proteins were found to be associated with tumor bone metastasis, tumors, and bone-related diseases. In the early stages of disease in model rats, these disease-associated proteins already existed. Previous studies have also directly proved the correctness of animal models and verified our results. Although these differential proteins are associated with many diseases and lack specificity alone, they may be better able to reflect bone metastases when combined. This indicates that the urine proteome can respond to diseases early and sensitively and is an ideal source of early biomarkers for tumor bone metastasis.

### 2.5 Results of GO analysis of the differentially proteins

For the differential proteins screened at different time points, we used the online database DAVID for functional analysis to obtain the following four results: biological processes (see Table S1), cellular components (see Table S2), molecular functions (Table 2) and pathways (Table 3).

**Table 2:**
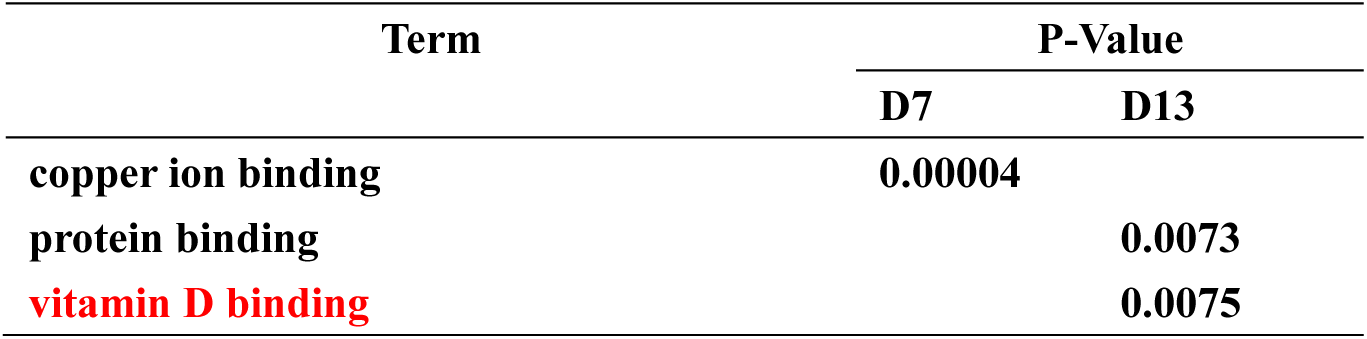
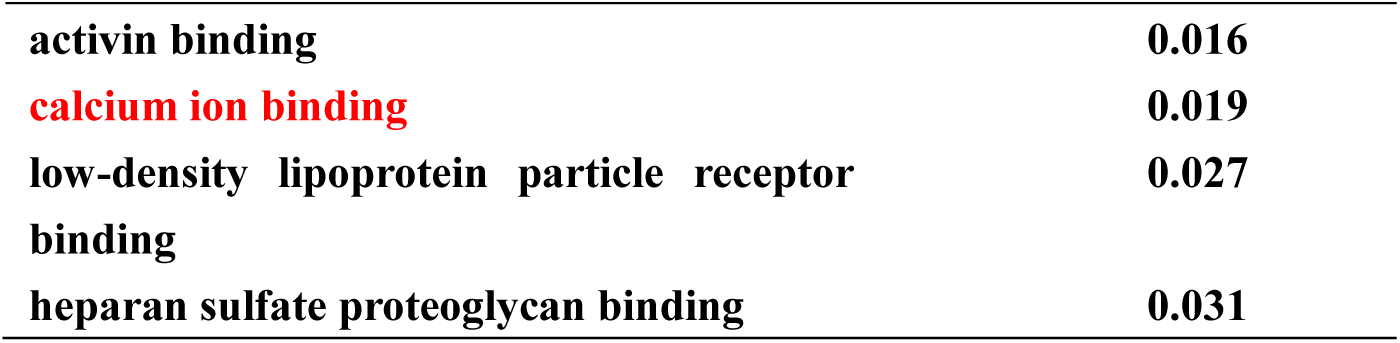
Results of the molecular function analysis of differential proteins

| Term | P-Value |  |
| --- | --- | --- |
|  | D7 | D13 |
| **copper ion binding** | **0.00004** |  |
| **protein binding** |  | **0.0073** |
| **vitamin D binding** |  | **0.0075** |

|  |  |
| --- | --- |
| activin binding | 0.016 |
| calcium ion binding | 0.019 |
| low-density lipoprotein particle receptor binding | 0.027 |
| heparan sulfate proteoglycan binding | 0.031 |

**Table 3:** Results of the pathway analysis for the differential proteins

| Term | P-Value |  |  |  |
| --- | --- | --- | --- | --- |
|  | D3 | D5 | D7 | D13 |
| Lysosome | 0.027 |  |  |  |
| Viral myocarditis |  | 0.0035 |  |  |
| Cell adhesion molecules (CAMs) |  | 0.012 |  |  |
| Mineral absorption |  |  | 0.0014 |  |
| Complement and coagulation cascades |  |  | 0.0044 |  |
| Folate biosynthesis |  |  |  | 0.023 |

We drew a graph of the biological processes of the differential proteins (Fig. 5), and we found that the immune response, such as the acute phase response, the adaptive immune response and the innate immune response, occurred on the third day after the tumor cells were implanted. These biological processes still existed on the seventh day (there were no obvious immune responses on days 5 and 13 that may be due to fewer differential proteins). In summary, we can see that the changes in the body against the tumor cells can be reflected early and with a high sensitivity in the urine. In this model, we also observed some ion reactions, such as the reaction to iron ions on the third and seventh days and the reaction to copper ions on the fifth and seventh days. On the 13th day, biological processes, such as wound healing and body development, occurred. These processes may all be related to tumor progression and may require our research to validate.

**Figure 5:**
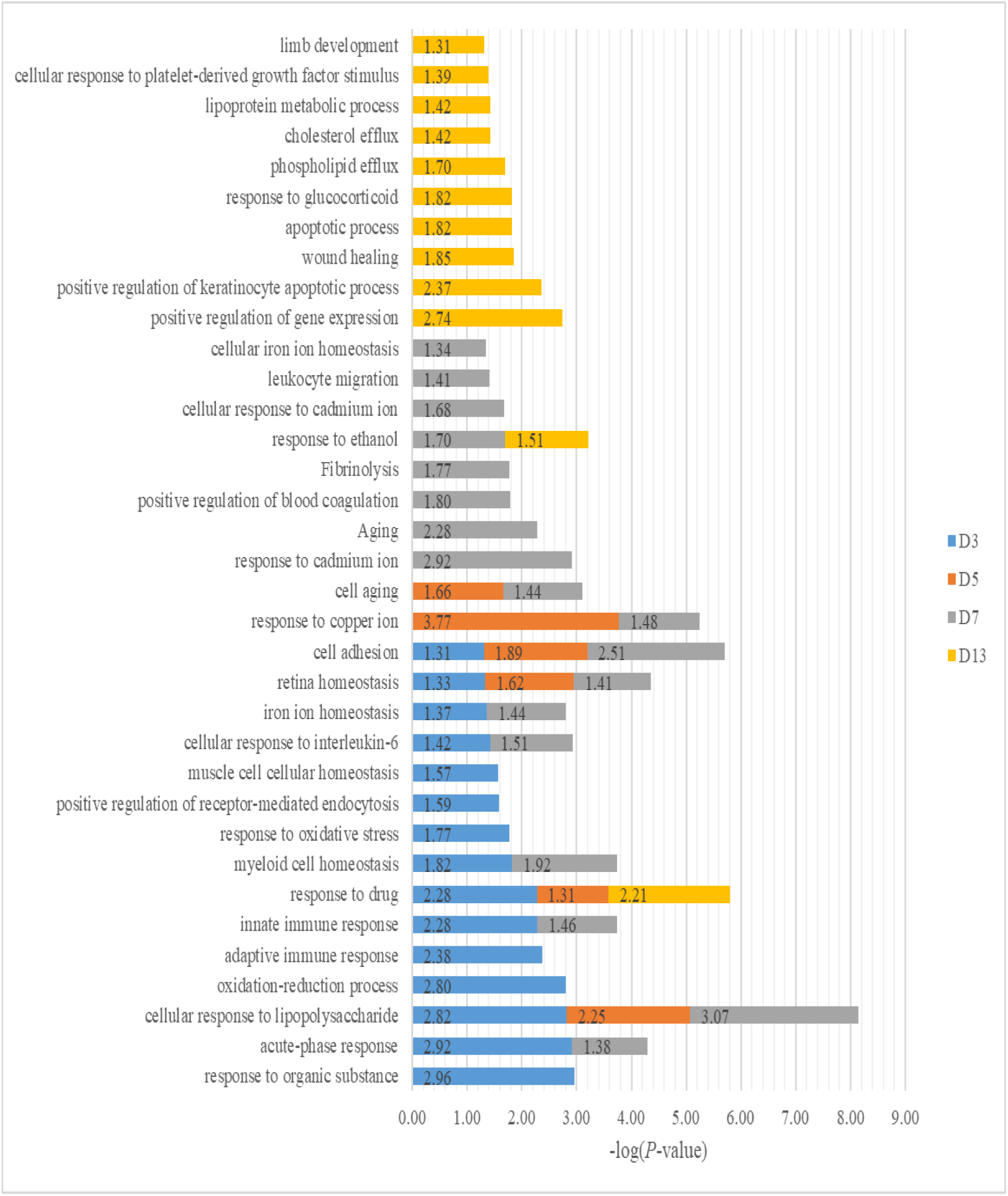
Results of the biological process analysis of differential proteins (D3, D5, D7, and D13: Walker 256 cells implanted on days 3, 5, 7, and 13)

We performed molecular functions (Table 2) and pathway analysis on differential proteins (Table 3). The molecular functions showed copper ion binding (marked in red) on the 7th day, which coincided with the biological process on the 7th day. On day 13, vitamin D binding and calcium ion binding (red part) appeared, and these functions may be related to bone remodeling and destruction. In the biological pathway, mineral absorption (marked in red) appeared on the 7th day, which is also likely to be related to bone destruction and reconstruction.

From the results of the above GO analysis, we can see that the urine proteome can reflect the early changes produced by the body after implanting the tumor cells with high sensitivity. In the later stages of the tumor, the reactions and pathways associated with bone metastases can also be seen in the urine. All of the above findings indicate that our model can reflect disease progression and that urine is an ideal source for finding the early markers of tumor bone metastasis.

### 2.6 Comparison of urinary proteins in different tumor models

The differential urinary proteins of five Walker 256 tumor models (liver tumor model, lung metastasis model, intracerebral tumor model, subcutaneous tumor model, and implanted bone cancer model) at all time points were compared and are shown in Fig. 6.

**Figure 6:**
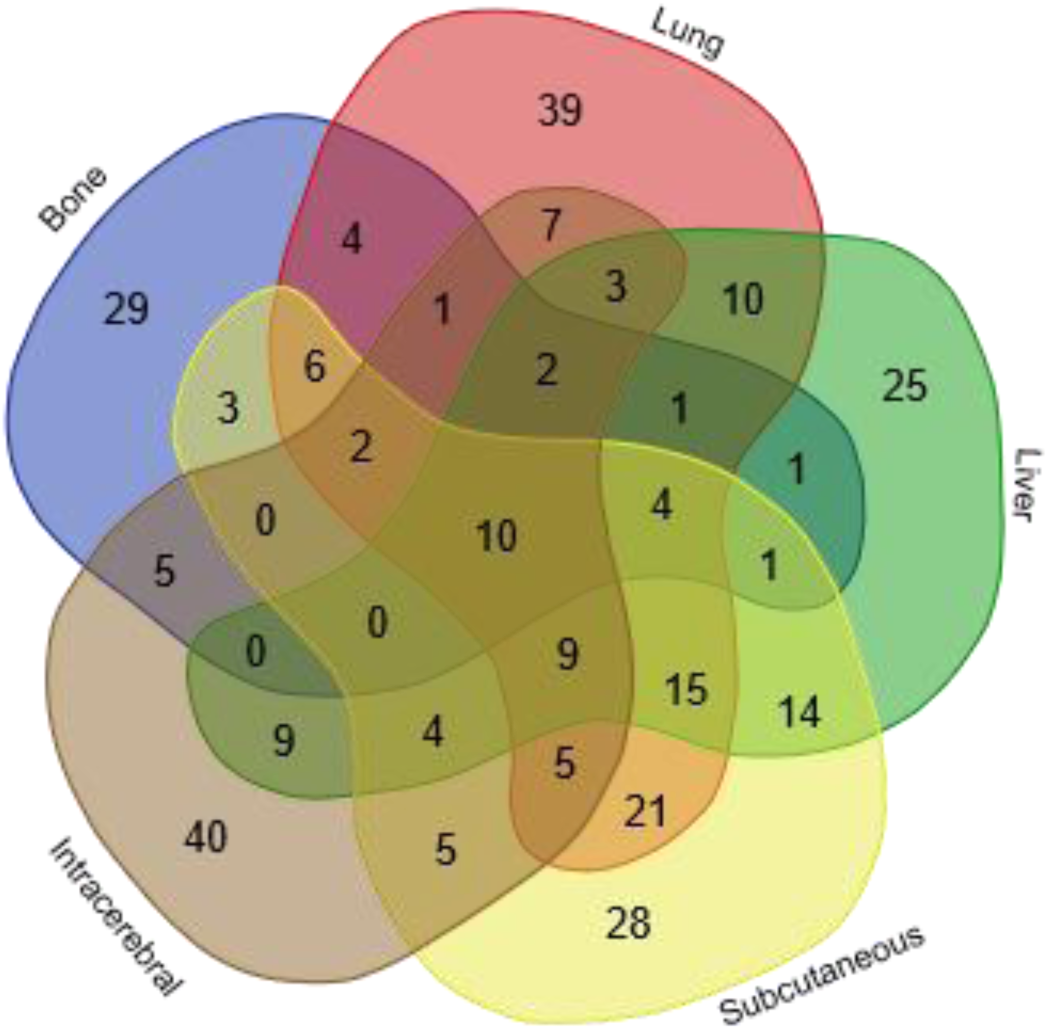
The overlapping differential proteins in the urine samples of the five different Walker 256 tumor models. (**Bone**: The differential proteins of the W Walker 256 256 implanted bone cancer model; **Lung**: The differential proteins of the Walker 256 lung metastasis model; **Liver**: The differential proteins of the Walker 256 liver tumor model; **Intracerebral**: The differential proteins of the Walker 256 intracerebral tumor model; **Subcutaneous**: The differential proteins of the Walker 256 Subcutaneous tumor model).

The results show that the changes produced by transplanting the same tumor cells into different body parts of rats could be distinguished in the urine. In Fig. 6, each model has unique differential proteins. The 29, 25, 39, 40, and 28 unique differential proteins were identified in the implanted bone cancer model, liver tumor model, lung metastasis model, intracerebral tumor model, and subcutaneous model, respectively.

One of the 29 proteins has been reported to be associated with bone metastasis. The protein is Follistatin. In prostate cancer patients, the serum concentrations of Follistatin have been shown to significantly correlate with the presence of bone metastases (P = 0.0005) [29]. Five of the 29 proteins have been reported to be associated with orthopedic disease: 1) The hepcidin gene showed a significant correlation with the RA disease activity parameters [30]. 2) Thioredoxin was a biomarker for oxidative stress in patients with rheumatoid arthritis [31]. 3) Metallothionein-1 (MT-1) was upregulated in RA and may prove to be a potential therapeutic target for RA autoimmune diseases [32]. 4) Mucosal addressin cell adhesion molecule 1 was strongly expressed in the synovium of osteoarthritis patients [33]. 5) The serum levels of the apolipoprotein C-I, C-III, and an N-terminal truncated form of transthyretin differed significantly between OA progressors and nonprogressors and are expected to be prognostic biomarkers for knee OA [34].

Among the overlapping proteins of these five models, it can be found that (1) 10 proteins can be detected in all models; and (2) most of the overlapping proteins reappear in more than two models in different combinations (Fig. 6). We think the reason for the appearance of these common proteins may be due to the same cell type that was injected into all models. (3) The different combinations of common proteins are also important for diagnosis. Because a single biomarker is difficult to diagnose, it is difficult to diagnose the type of tumor. The biomarker panel is more accurate and reliable. In conclusion, the results of the comparison show that the growth of tumors in different organs has both similarities and individual differences. Urinary proteins may differentiate the same tumor cells that grow in different organs.

---

**Table S1:**
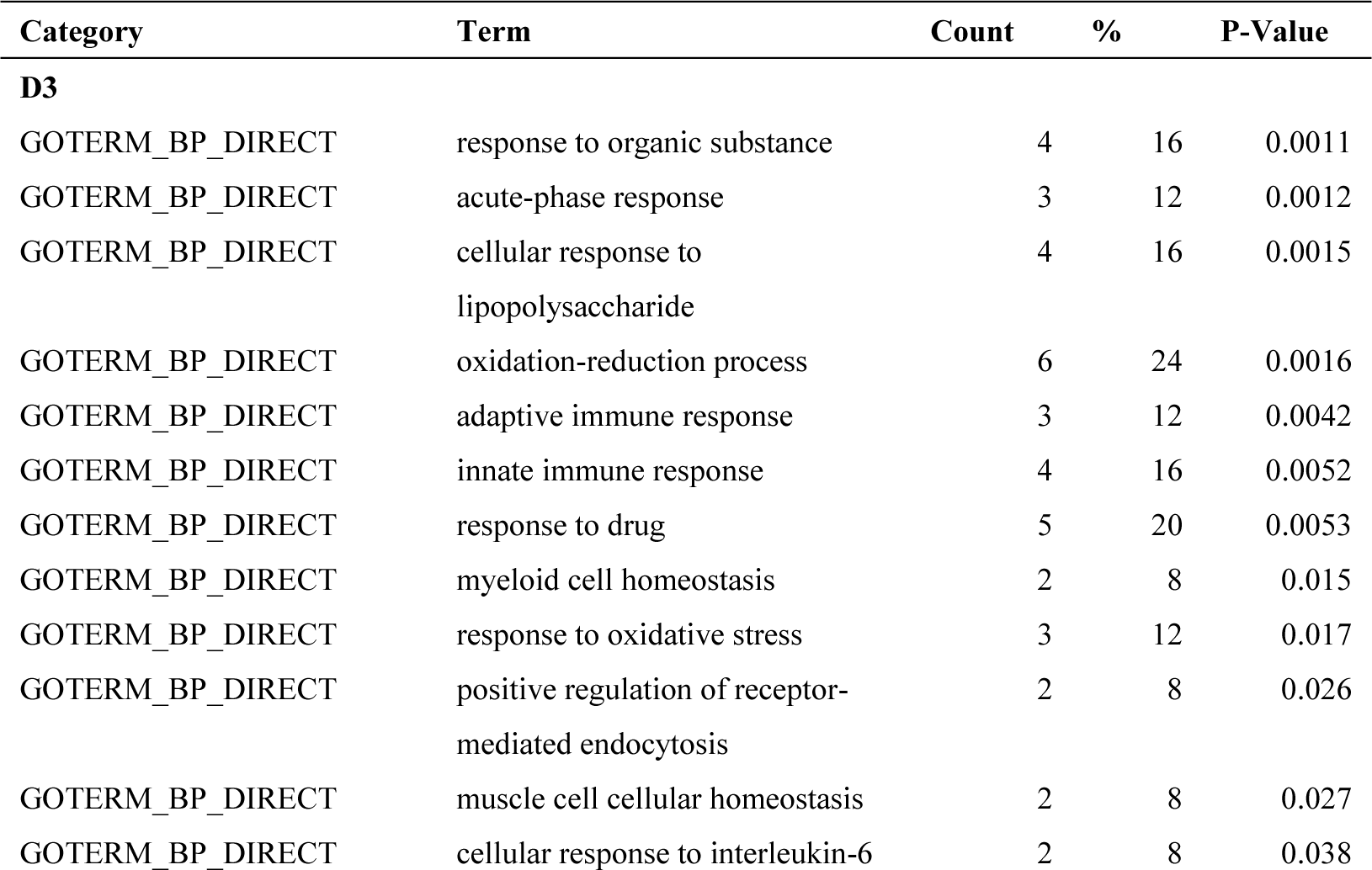
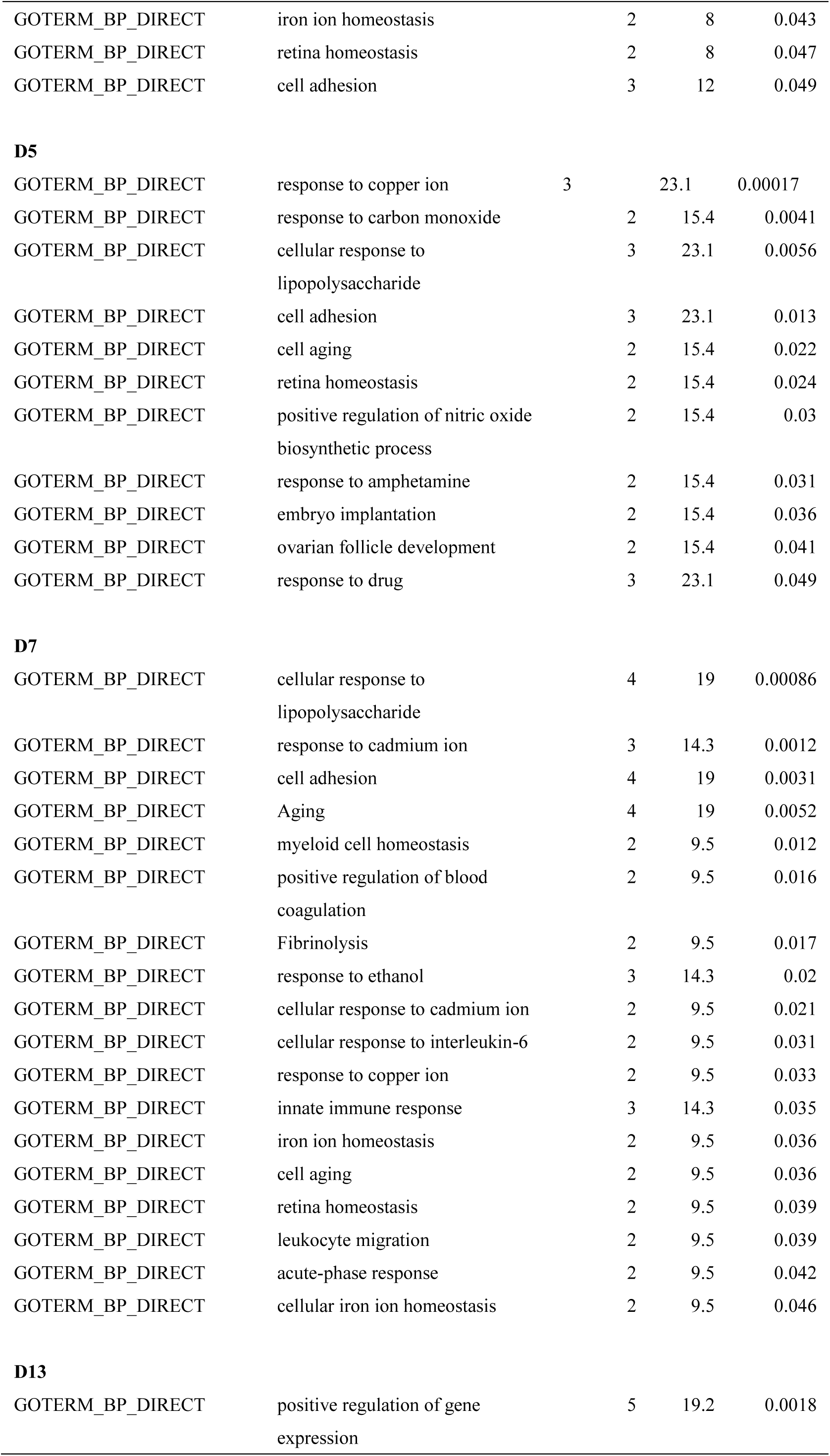
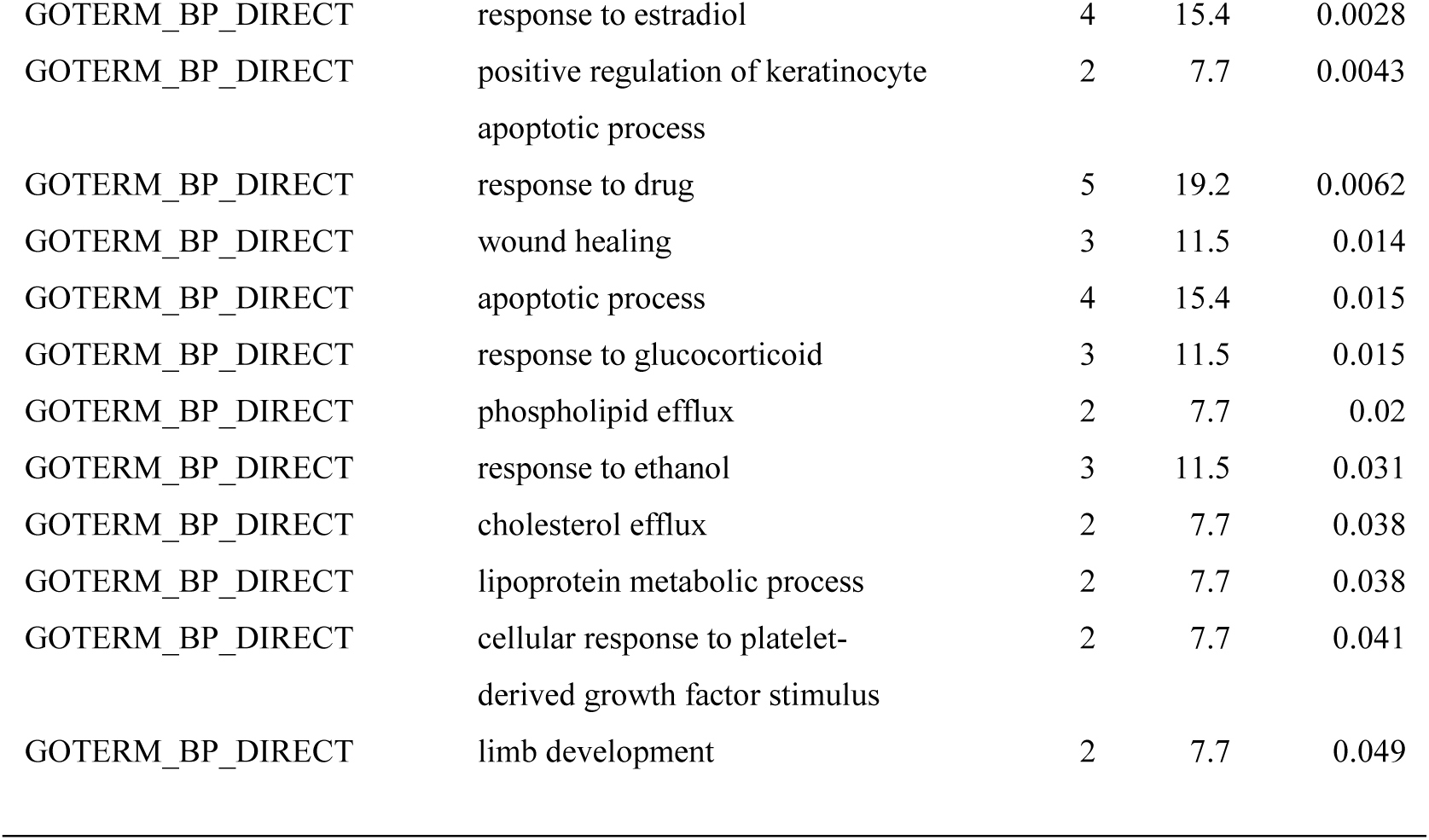
Biological processes of differential proteins.

| Category | Term | Count | % | P-Value |
| --- | --- | --- | --- | --- |
| **D3** |  |  |  |  |
| GOTERM_BP_DIRECT | response to organic substance | 4 | 16 | 0.0011 |
| GOTERM_BP_DIRECT | acute-phase response | 3 | 12 | 0.0012 |
| GOTERM_BP_DIRECT | cellular response to lipopolysaccharide | 4 | 16 | 0.0015 |
| GOTERM_BP_DIRECT | oxidation-reduction process | 6 | 24 | 0.0016 |
| GOTERM_BP_DIRECT | adaptive immune response | 3 | 12 | 0.0042 |
| GOTERM_BP_DIRECT | innate immune response | 4 | 16 | 0.0052 |
| GOTERM_BP_DIRECT | response to drug | 5 | 20 | 0.0053 |
| GOTERM_BP_DIRECT | myeloid cell homeostasis | 2 | 8 | 0.015 |
| GOTERM_BP_DIRECT | response to oxidative stress | 3 | 12 | 0.017 |
| GOTERM_BP_DIRECT | positive regulation of receptor-mediated endocytosis | 2 | 8 | 0.026 |
| GOTERM_BP_DIRECT | muscle cell cellular homeostasis | 2 | 8 | 0.027 |
| GOTERM_BP_DIRECT | cellular response to interleukin-6 | 2 | 8 | 0.038 |
| GOTERM_BP_DIRECT | iron ion homeostasis | 2 | 8 | 0.043 |
| GOTERM_BP_DIRECT | retina homeostasis | 2 | 8 | 0.047 |
| GOTERM_BP_DIRECT | cell adhesion | 3 | 12 | 0.049 |
| **D5** |  |  |  |  |
| GOTERM_BP_DIRECT | response to copper ion | 3 | 23.1 | 0.00017 |
| GOTERM_BP_DIRECT | response to carbon monoxide | 2 | 15.4 | 0.0041 |
| GOTERM_BP_DIRECT | cellular response to lipopolysaccharide | 3 | 23.1 | 0.0056 |
| GOTERM_BP_DIRECT | cell adhesion | 3 | 23.1 | 0.013 |
| GOTERM_BP_DIRECT | cell aging | 2 | 15.4 | 0.022 |
| GOTERM_BP_DIRECT | retina homeostasis | 2 | 15.4 | 0.024 |
| GOTERM_BP_DIRECT | positive regulation of nitric oxide biosynthetic process | 2 | 15.4 | 0.03 |
| GOTERM_BP_DIRECT | response to amphetamine | 2 | 15.4 | 0.031 |
| GOTERM_BP_DIRECT | embryo implantation | 2 | 15.4 | 0.036 |
| GOTERM_BP_DIRECT | ovarian follicle development | 2 | 15.4 | 0.041 |
| GOTERM_BP_DIRECT | response to drug | 3 | 23.1 | 0.049 |
| **D7** |  |  |  |  |
| GOTERM_BP_DIRECT | cellular response to lipopolysaccharide | 4 | 19 | 0.00086 |
| GOTERM_BP_DIRECT | response to cadmium ion | 3 | 14.3 | 0.0012 |
| GOTERM_BP_DIRECT | cell adhesion | 4 | 19 | 0.0031 |
| GOTERM_BP_DIRECT | Aging | 4 | 19 | 0.0052 |
| GOTERM_BP_DIRECT | myeloid cell homeostasis | 2 | 9.5 | 0.012 |
| GOTERM_BP_DIRECT | positive regulation of blood coagulation | 2 | 9.5 | 0.016 |
| GOTERM_BP_DIRECT | Fibrinolysis | 2 | 9.5 | 0.017 |
| GOTERM_BP_DIRECT | response to ethanol | 3 | 14.3 | 0.02 |
| GOTERM_BP_DIRECT | cellular response to cadmium ion | 2 | 9.5 | 0.021 |
| GOTERM_BP_DIRECT | cellular response to interleukin-6 | 2 | 9.5 | 0.031 |
| GOTERM_BP_DIRECT | response to copper ion | 2 | 9.5 | 0.033 |
| GOTERM_BP_DIRECT | innate immune response | 3 | 14.3 | 0.035 |
| GOTERM_BP_DIRECT | iron ion homeostasis | 2 | 9.5 | 0.036 |
| GOTERM_BP_DIRECT | cell aging | 2 | 9.5 | 0.036 |
| GOTERM_BP_DIRECT | retina homeostasis | 2 | 9.5 | 0.039 |
| GOTERM_BP_DIRECT | leukocyte migration | 2 | 9.5 | 0.039 |
| GOTERM_BP_DIRECT | acute-phase response | 2 | 9.5 | 0.042 |
| GOTERM_BP_DIRECT | cellular iron ion homeostasis | 2 | 9.5 | 0.046 |
| **D13** |  |  |  |  |
| GOTERM_BP_DIRECT | positive regulation of gene expression | 5 | 19.2 | 0.0018 |
| GOTERM_BP_DIRECT | response to estradiol | 4 | 15.4 | 0.0028 |
| GOTERM_BP_DIRECT | positive regulation of keratinocyte apoptotic process | 2 | 7.7 | 0.0043 |
| GOTERM_BP_DIRECT | response to drug | 5 | 19.2 | 0.0062 |
| GOTERM_BP_DIRECT | wound healing | 3 | 11.5 | 0.014 |
| GOTERM_BP_DIRECT | apoptotic process | 4 | 15.4 | 0.015 |
| GOTERM_BP_DIRECT | response to glucocorticoid | 3 | 11.5 | 0.015 |
| GOTERM_BP_DIRECT | phospholipid efflux | 2 | 7.7 | 0.02 |
| GOTERM_BP_DIRECT | response to ethanol | 3 | 11.5 | 0.031 |
| GOTERM_BP_DIRECT | cholesterol efflux | 2 | 7.7 | 0.038 |
| GOTERM_BP_DIRECT | lipoprotein metabolic process | 2 | 7.7 | 0.038 |
| GOTERM_BP_DIRECT | cellular response to platelet-derived growth factor stimulus | 2 | 7.7 | 0.041 |
| GOTERM_BP_DIRECT | limb development | 2 | 7.7 | 0.049 |

**Table S2:** Cellular components of differential proteins

| Category | Term | Count | % | P-Value |
| --- | --- | --- | --- | --- |
| **D3** |  |  |  |  |
| GOTERM_CC_DIRECT | extracellular space | 14 | 56 | 1.3E-09 |
| GOTERM_CC_DIRECT | extracellular exosome | 15 | 60 | 6.8E-07 |
| GOTERM_CC_DIRECT | Lysosome | 4 | 16 | 0.0037 |
| GOTERM_CC_DIRECT | neuronal cell body | 5 | 20 | 0.0048 |
| GOTERM_CC_DIRECT | external side of plasma membrane | 4 | 16 | 0.0048 |
| GOTERM_CC_DIRECT | blood microparticle | 3 | 12 | 0.01 |
| GOTERM_CC_DIRECT | membrane raft | 3 | 12 | 0.046 |
| **D5** |  |  |  |  |
| GOTERM_CC_DIRECT | extracellular exosome | 9 | 69.2 | 0.00005 |
| GOTERM_CC_DIRECT | extracellular space | 5 | 38.5 | 0.0079 |
| GOTERM_CC_DIRECT | external side of plasma membrane | 3 | 23.1 | 0.012 |
| GOTERM_CC_DIRECT | MHC class I protein complex | 2 | 15.4 | 0.015 |
| GOTERM_CC_DIRECT | mitochondrial intermembrane space | 2 | 15.4 | 0.045 |
| **D7** |  |  |  |  |
| GOTERM_CC_DIRECT | extracellular space | 12 | 57.1 | 2.1E-08 |
| GOTERM_CC_DIRECT | extracellular exosome | 13 | 61.9 | 2.9E-06 |
| GOTERM_CC_DIRECT | external side of plasma membrane | 4 | 19 | 0.0028 |
| GOTERM_CC_DIRECT | blood microparticle | 3 | 14.3 | 0.0071 |
| GOTERM_CC_DIRECT | extracellular matrix | 3 | 14.3 | 0.03 |
| GOTERM_CC_DIRECT | extracellular region | 4 | 19 | 0.04 |
| **D13** |  |  |  |  |
| GOTERM_CC_DIRECT | extracellular exosome | 23 | 88.5 | 3.5E-16 |
| GOTERM_CC_DIRECT | extracellular space | 13 | 50 | 3.4E-08 |
| GOTERM_CC_DIRECT | blood microparticle | 5 | 19.2 | 0.000018 |
| GOTERM_CC_DIRECT | extracellular region | 8 | 30.8 | 0.000032 |
| GOTERM_CC_DIRECT | Lysosome | 4 | 15.4 | 0.0042 |
| GOTERM_CC_DIRECT | Cytosol | 7 | 26.9 | 0.017 |
| GOTERM_CC_DIRECT | high-density lipoprotein particle | 2 | 7.7 | 0.023 |
| GOTERM_CC_DIRECT | very-low-density lipoprotein particle | 2 | 7.7 | 0.023 |
| GOTERM_CC_DIRECT | myelin sheath | 3 | 11.5 | 0.027 |
| GOTERM_CC_DIRECT | cell surface | 4 | 15.4 | 0.048 |

